# The tortoise and the hare: interactions between reinforcement learning and working memory

**DOI:** 10.1101/234724

**Authors:** Anne G.E. Collins

**Affiliations:** Department of Psychology, Helen Wills Neuroscience Institute, UC Berkeley.

## Abstract

Learning to make rewarding choices in response to stimuli depends on a slow but steady process, reinforcement learning, and a fast and flexible, but capacity limited process, working memory. Using both systems in parallel, with their contributions weighted based on performance, should allow us to leverage the best of each system: rapid early learning, supplemented by long term robust acquisition. However, this assumes that using one process does not interfere with the other. We use computational modeling to investigate the interactions between the two processes in a behavioral experiment, and show that working memory interferes with reinforcement learning. Previous research showed that neural representations of reward prediction errors, a key marker of reinforcement learning, were blunted when working memory was used for learning. We thus predicted that arbitrating in favor of working memory to learn faster in simple problems would weaken the reinforcement learning process. We tested this by measuring performance in a delayed testing phase where the use of working memory was impossible, and thus subject choices depended on reinforcement learning. Counter-intuitively, but confirming our predictions, we observed that associations learned most easily were retained worse than associations learned slower: using working memory to learn quickly came at the cost of long-term retention. Computational modeling confirmed that this could only be accounted for by working memory interference in reinforcement learning computations. These results further our understanding of how multiple systems contribute in parallel to human learning, and may have important applications for education and computational psychiatry.

## Introduction

When facing a challenge (such as responding to a natural disaster), we often need to pursue two solutions in parallel: emergency measures for the immediate future, and carefully thought-out long-term plans. These measures make different trade-offs between speed and efficiency, and neither is better than the other in absolute. Allocating finite resources to multiple strategies that involve such different trade-offs can mitigate their limitations and make use of their benefits. Recent work shows that human learning, even in very simple environments, follows this principle: it involves multiple parallel neurocognitive systems that arbitrate differently between immediate, effortful efficacy, and slower, long-term robustness (Bornstein, Khaw, Shohamy, & Daw, 2017; Collins & Frank, 2012; Daw, Gershman, Seymour, Dayan, & Dolan, 2011; R. Poldrack et al., 2001). In this article, we present a new experiment to computationally characterize how these systems work together to accomplish short- and long-term learning.

We focus on two well-defined systems, reinforcement learning (RL) and working memory (WM). WM enables fast and accurate single-trial learning of any kind of information, with two limitations – a capacity or resource limit, and a temporal limit, such that we can only accurately remember a small amount of information for a short time, after which we may forget (Baddeley, 2012). RL, in contrast, enables slower but robust reward-based learning of the value of choices (Dayan & Daw, 2008). Separable (yet partially overlapping) brain networks support these two systems: WM is centrally dependent on prefrontal cortex (Cools, 2011; Miller & Cohen, 2001), while RL relies on dopaminergic signaling in the striatum (Frank & O’Reilly, 2006; Montague, Dayan, & Sejnowski, 1996; Schultz, 2013; Tai, Lee, Benavidez, Bonci, & Wilbrecht, 2012). We previously developed a simple experimental protocol to show that both systems are used in parallel for instrumental learning (Collins, Ciullo, Frank, & Badre, 2017; Collins & Frank, 2012). In this protocol, participants used deterministic feedback to learn the correct action to pick in response to a stimulus; critically, in different blocks, they needed to learn associations for a different number of stimuli in a new set. This *set size* manipulation allowed us to disentangle the contributions of WM to learning from those of RL, because WM is capacity-limited (Cowan, 2010), while RL is not. In particular, this manipulation was essential in identifying which of the systems was causing learning dysfunction in clinical populations (Collins, Albrecht, Waltz, Gold, & Frank, 2017; Collins, Brown, Gold, Waltz, & Frank, 2014).

However, how the two systems interact is still poorly understood. Previous models assumed that the RL and WM systems compete for choice, but otherwise learn independently of each other. In particular, if RL was fully independent from WM, values learned through RL should only depend on reward history, and be independent of set size. Recent evidence shows that this may not be the case, and that WM may instead interfere in RL computations. Indeed, in an fMRI study, we showed that RL signals were blunted in low set sizes, when working memory was sufficient to learn stimulus-response associations (Collins, Ciullo, et al., 2017). These results were confirmed in an EEG study (Collins & Frank, submitted), where we additionally found that trial-by-trial markers of WM use predicted weaker RL learning signals. Last, we found behavioral evidence that the value of different items was better retained when learned under high load than under low load (Collins, Albrecht, Waltz, Gold, & Frank, 2017), again hinting at an interference of WM with RL computations. Together, these results hint at a mechanism by which RL learning is weakened in low set sizes, when working memory is most successful. We hypothesize that successful WM use (in particular in low set sizes) may interfere with RL, either by inhibiting learning, or, more consistently with our most recent EEG findings, by communicating to the RL system its quickly acquired expectations and thus making positive reward prediction error signals less strong.

In this project, we further investigate the nature of interactions between WM and RL during learning. Specifically, we propose an improvement to our previous protocol that allows us to 1) characterize short-term learning vs. long-term learning and retention, 2) investigate how WM use impacts both forms of learning, and 3) computationally characterize the complete, integrated, interactive learning process. In the new protocol (Fig. 1A), the learning phase includes multiple blocks of one of two set sizes – low (3) and high (6); after a short, irrelevant task to provide a delay, the experimental protocol ends with a surprise testing phase in extinction to assess the retention of the associations acquired during the learning phase. This testing phase probes how well participants remember the correct choice for all the stimuli they learned previously. We hypothesize that WM plays no direct role in testing phase choices (Fig. 1B) because, in the absence of feedback, no new information is available in the testing phase: participants decide based on experience acquired more than 10 minutes prior, for 54 different stimulus-action associations, both beyond the extent of working memory maintenance. Thus, the testing phase serves as a purer marker of RL function than could be obtained with the original design (Collins & Frank, 2012), allowing us to observe how RL-learned associations depend on the learning context (high/low set size); and thus to investigate the interaction of WM and RL.

**Figure 1:**
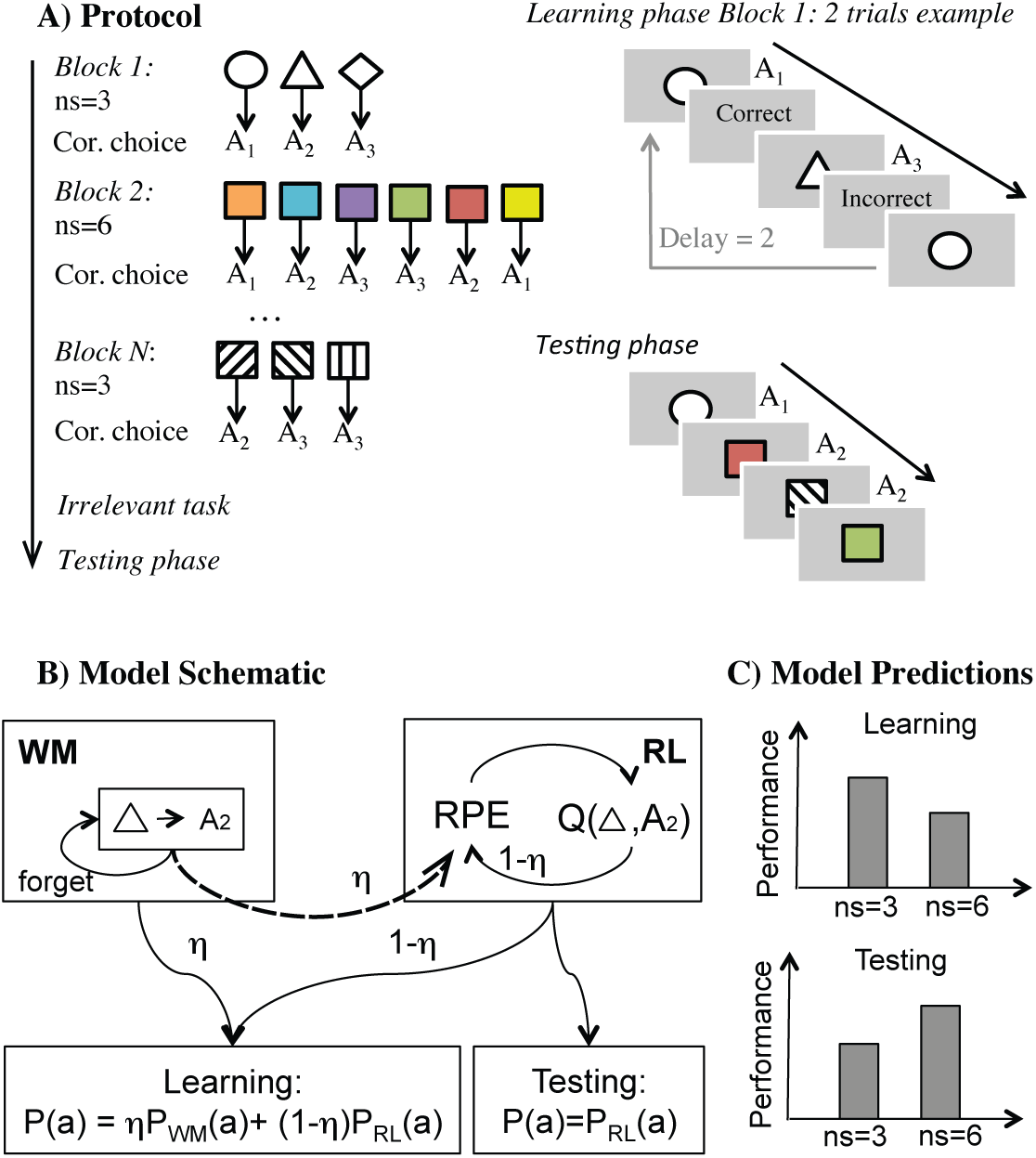
**A) Experimental protocol**. Left: the experiment includes a learning phase with 14 learning blocks of set size *ns*=*3* or *6*; followed by a short irrelevant task and a surprise testing phase. Right: examples of learning phase and testing phase trials. **B) Model schematic.** We assume that WM and RL both contribute competitively to choice during learning; the weight *η* of WM vs. RL for choice depends on the capacity and set size. WM stores exact information but may forget; RL learns from reward prediction errors (RPE) the value of selecting an action for a given stimulus (e.g. the triangle). We additionally hypothesize that WM influences RL computations (dashed arrow) by contributing expectations to the computation of the RPE. Testing phase does not involve WM contributions, but only RL. **C) Model predictions (schematic):** if *η*(*ns*=*3*)<*η*(*ns*=*6*), the model predicts worse performance under high load during learning; however items learned under high load are predicted to be better retained during testing, because WM interferes less with RL.

Based on our previous imaging results showing blunted RL signals in low set sizes, we made the counter-intuitive prediction that retention of associations learned under low set sizes would be worse than in high set sizes (Fig. 1C). Furthermore, we predicted that a model incorporating interference of WM in RL computations (Fig. 1B, methods) would capture behavior better than models assuming either single systems or independent WM and RL modules competing for choice.

## Methods

### Subjects

We first tested 49 University of California, Berkeley undergraduate students (31 female; ages 18-40, mean 20.5). To replicate the results obtained with this first sample, we then tested a second, independent sample, which included 42 University of California, Berkeley undergraduate students (26 female; ages 18-29, mean 20.7). Because the effects in the second sample fully replicated the results obtained with the first sample, we report here the results with participants pooled together. We excluded 6 participants based on poor overall performance in the task indicating lack of involvement (less than 75% average accuracy at asymptotic performance). The final sample included N = 85 participants. All participants provided written informed consent, and the Committee for the Protection of Human Subjects at the University of California, Berkeley, approved the research.

### Experimental protocol

#### General

The methods are modified from previous published versions of this experimental protocol (Collins, Albrecht, et al., 2017; Collins et al., 2014; Collins, Ciullo, et al., 2017; Collins & Frank, 2012, submitted). Subjects performed a learning experiment in which they used reinforcement feedback (“+1” or “0”) to figure out the correct key to press in response to several visual stimuli (Fig. 1A). The experiment was separated into three phases: a learning phase comprising multiple independent learning blocks (mean duration 26 min, range [22-34]); an unrelated memory task (mean duration 11 min, range [10-16]; and a surprise testing phase assessing what was retained from the learning phase (mean duration 7 min, range [6-8]).

#### Learning phase

The learning phase was separated into 14 blocks, with a new set of visual stimuli of set size *ns* in each block, with *ns*=*3* or *ns*=*6*. Each block included 12-14 presentations of each visual stimuli in a pseudo-randomly interleaved manner (controlling for a uniform distribution of delay between two successive presentations of the same stimulus within [*1:2*ns*] trials, and the number of presentations of each stimulus), for a total of *ns*\**13* trials. At each trial, a stimulus was presented centrally on a black background. Subjects had up to 1.5 seconds to answer by pressing one of three keys with their right hand. Key press was followed by visual feedback presentation for 0.5 seconds, followed by a fixation of 0.5 seconds before the onset of the next trial. For each image, the correct key press was the same for the entire block. Pressing this key led to a truthful “+1” feedback, while pressing any other key led to a truthful “0” feedback. Failure to answer within 1.5 seconds was indicated by a “no valid answer” message. Stimuli in a given block were all from a single category (e.g. colors, fruits, animals), and did not repeat across blocks.

We varied the set size *ns* across blocks: out of 14 blocks, 8 had set size *ns*=*3* and 6 set size *ns*=*6*. The first and last block of the learning phase were of set size *ns*=*3*. We have shown in previous research that varying set size provides a way to investigate the contributions of capacity- and resource-limited working memory to reinforcement learning.

#### N-back

Following the learning phase, participants performed a classic visual N-back task (N = 2-5). The purpose of this phase was to provide a time buffer between the learning phase and the surprise testing phase. We do not analyze the N-back performance here.

#### Testing phase

At the end of the N-back task, participants were informed that they would be tested on what they had learned during the first phase of the experiment. The testing phase included a single block where all stimuli from the learning phase (excepting block 1 and block N to limit recency and primacy effects – thus a total of 54 stimuli: 36 learned in set size 6, and 18 learned in set size 3 blocks) were presented 4 times each, for a total of 216 trials. The order was pseudo-randomized to ensure that each stimulus was presented once in each quarter of the test phase. Each trial was identical to the learning phase, except that no feedback was presented so that participants could not learn during this phase.

### Model free analysis

#### Learning phase

Learning curves were constructed by computing the proportion of correct answers to all stimuli of each set size as a function of their iteration number. Trials with missing answers were excluded. Reaction time learning curves were limited to correct choices. Asymptotic performance/reaction times were assessed over the last 6 presentations of each stimulus.

We also used a multiple logistic/linear regression analysis to predict correct choices/reaction times in the learning phase. The main predictor was the “reward” predictor (*#R*), which indicated the number of previous correct choices for a given stimulus and was expected to capture the effect of reward history on learning (Collins & Frank, 2012). Predictors expected to capture working memory contributions were 1) the set size *ns* of the block in which a stimulus was learned and 2) the delay, or number of trials since the subject last selected the correct action for the current stimulus (Fig. 1A). Last, we also included the block number to investigate whether exposure to the learning phase modified performance. All regressors were z-scored prior to entering into the regression model. For visualization purposes in figures 2 and 5, weights are sigmoid transformed and scaled to [-1,1] (keeping 0 as the no effect value). Specifically, the transformation is *β***←***(2/(1+exp(-β)))-1*.

**Figure 2:**
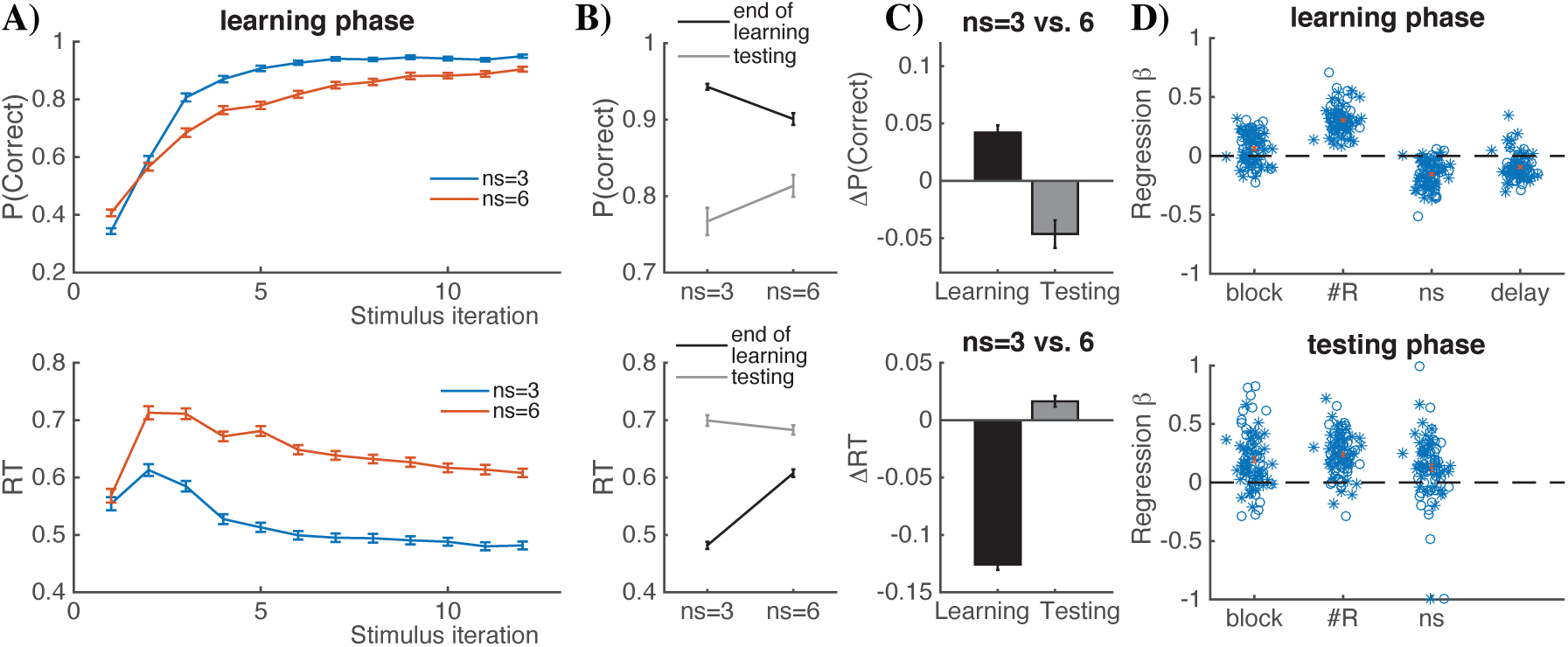
Behavioral results show opposite effect of load in learning and retention. **A-top)** Proportion of correct trials as a function of stimulus iteration number, for set sizes *ns = 3/6.* **A-bottom)** Same for reaction times (RT). **B-top)** Proportion of correct trials at asymptotic learning performance (last 5 iterations of a stimulus - black line) and during test phase – grey line). **B-bottom)** RT’s for testing and asymptotic learning. **C)** Difference between low and high set sizes shows significant, opposite results for learning and test phase, in both performance (top) and RT’s (bottom). **D-top)** Transformed logistic regression weights from learning phase show expected effects of learning block and reward history (due to practice and reinforcement learning), as well as expected negative effects of set size and delay, characterizing working memory contributions. **D-bottom)** Transformed logistic regression weights from the test phase also show expected effects of learning block and reward history, but show better test phase performance with higher set size, contrary to learning phase. Open circles represent individual subjects from original experiment; stars represent individual subjects from the replication sample. The results are identical. Error bars indicate standard error of the mean.

**Figure 5:**
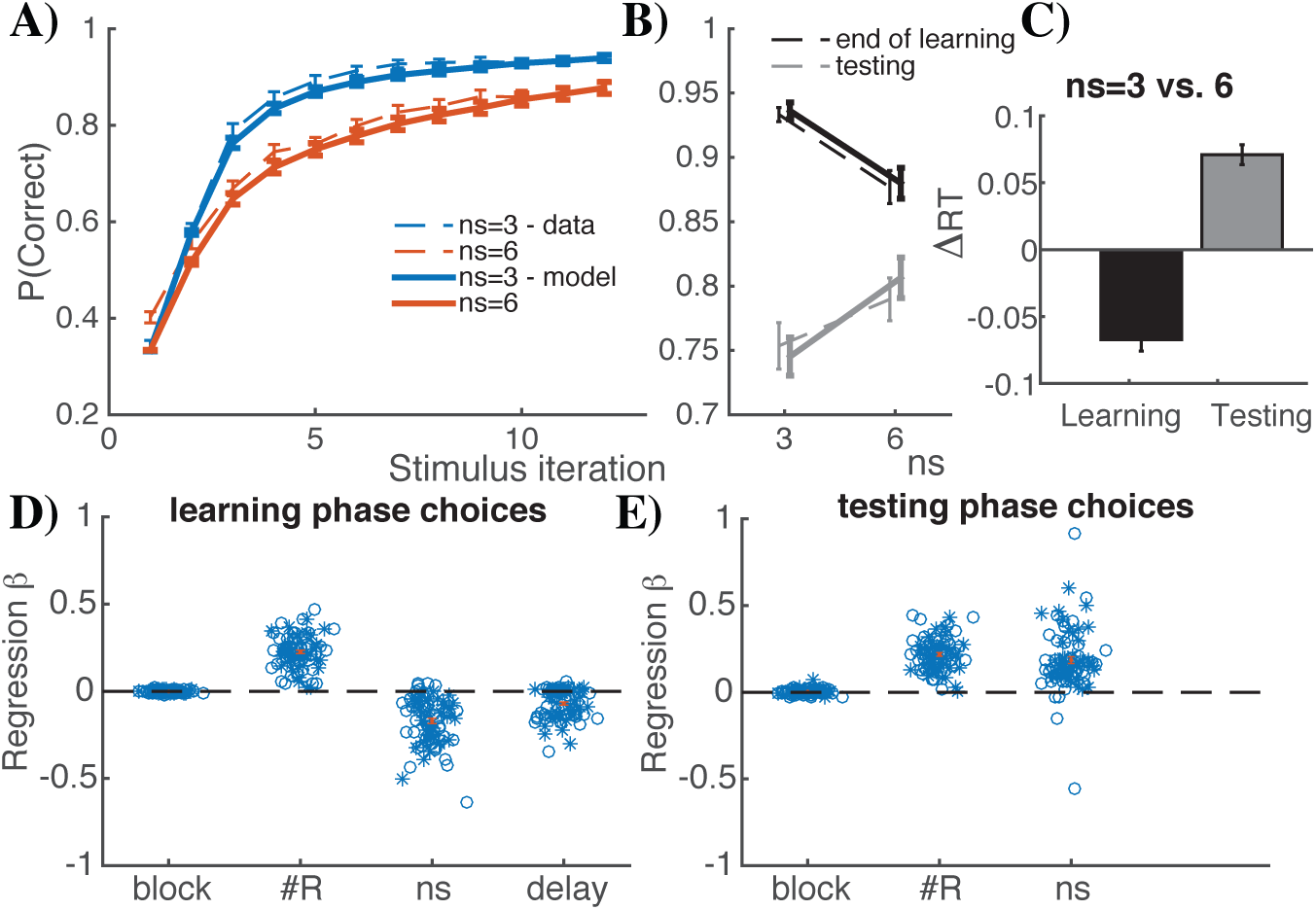
RLWMi validation. Model simulations with fit parameters. **A):** Model simulations reproduce learning curves. **B-C):** Model simulations reproduce the opposite effects of set size *ns* between learning and testing phases, for both choices and reaction times. **D-E):** Logistic regression analysis of the learning and testing phase choices capture behavioral effects (Fig. 2), including the opposite effects of set size on performance.

#### Testing phase

We computed the performance as the proportion of correct choices separately for stimuli that had been learned in blocks of set size *ns* = *3* or *ns* = *6*. We also used multiple logistic/linear regression analysis for the testing phase, with reward, set size, and block regressors. The reward regressor was the asymptotic performance for a given stimulus; and the set size and block regressors were defined as the set size and block number of the block during which a given stimulus had been learned in the learning phase.

#### Errors

To investigate the types of errors made during the testing phase we identified a set of stimuli that met the following criteria: 1) the participant had made at least one mistake for that stimulus during the learning phase, and 2) one of the two incorrect actions had been selected more often than the other during the learning phase. This allowed us to define for the testing phase a perseverative error as choosing the action that had been selected most, and an optimal error as choosing the erroneous action that had been selected least (this would correspond to best avoidance of previously unrewarding actions). We computed the proportion of perseverative vs. optimal errors separately for each set size.

#### Computational modeling

We simultaneously modeled the learning and testing phases based on our theory, as shown in Fig. 1C. The models assume that dynamically changing learned associations control learning phase policy, and that policies acquired at the end of the learning phase are used during the testing phase.

#### Models

We tested three families of models:

- ***RLs:*** Pure reinforcement learning models
- ***RLWM:*** mixture models with *independent* WM and RL; where both WM and RL contributed to learning, but only RL contributed to testing
- ***RLWMi:*** RLWM mixture models with *interacting* WM and RL; where both WM and RL contributed to learning, but only RL contributed to testing

### RL models

Computational models in the class of RL models are built off a classic formulation of RL models (see below), with two key parameters – learning rate *α* and softmax temperature *β*. We also investigate RL models that include various combinations of additional mechanisms that may help provide a better account of behavior, while remaining in the family of single process RL models. These additional mechanisms include undirected noise, forgetting, perseveration, initial bias, and dependence of learning rate on task conditions. We describe all mechanisms below.

#### Classic RL.

For each stimulus *s*, and action *a*, the expected reward *Q*(*s,a*) is learned as a function of reinforcement history. Specifically, the *Q* value for the selected action given the stimulus is updated upon observing each trial’s reward outcome r_t_ (1 for correct, 0 for incorrect) as a function of the prediction error between expected and observed reward at trial *t*:

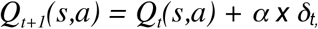

where *δ_t_*= *r_t_* − *Q_t_*(*s,a*) is the prediction error, and *α* is the learning rate. Choices are generated probabilistically with greater likelihood of selecting actions that have higher *Q*-values, using the softmax choice policy, which defines the probabilistic rule for choosing actions in response to a stimulus:

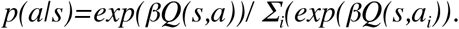

Here, *β* is an inverse temperature determining the degree with which differences in *Q*-values are translated into more deterministic choice, and the sum is over the three possible actions a_i_. Choice policy during the testing phase is identical to the policy at the end of the learning phase.

#### Undirected noise.

The softmax allows for stochasticity in choice, but stochasticity is more impactful when the value of each action is close to the values of the alternative actions. We also allow for “slips” of action (i.e., even when Q-value differences are large; also called “irreducible noise” or lapse rate). Given a model’s policy π = p(a|s), adding undirected noise consists in defining the new mixture choice policy:

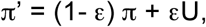

where U is the uniform random policy (U(a) = 1/n_A_, with number of actions n_A_=3), and the parameter 0 < ε < 1 controls the amount of noise (Collins & Frank, 2013; Collins & Koechlin, 2012; Guitart-Masip et al., 2012). Intuitively, this undirected noise captures a choice policy where with probability *1 − ε* the agent picks choices normally, but with probability *ε*, the agent lapses and acts randomly. (Nassar & Frank, 2016) showed that failing to take this irreducible noise into account can make model fits be unduly influenced by rare odd data points (e.g. that might arise from attentional lapses), and that this problem is remedied by using the hybrid softmax-ε-greedy choice function as used here.

#### Forgetting.

We allow for potential decay or forgetting in Q-values on each trial, additionally updating all Q-values at each trial, according to:

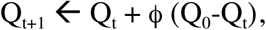

where 0 < ϕ < 1 is a decay parameter which at each trial pulls the estimates of values towards the initial value Q_0_ = 1/n_A_. This parameter allows us to capture forgetting during learning.

#### Perseveration.

To allow for potential neglect of negative, as opposed to positive feedback, we estimate a perseveration parameter *pers* such that for negative prediction errors (*δ* < 0), the learning rate *α* is reduced by *α* = *(1-pers)α*. Thus, values of *pers* near 1 indicate perseveration with complete neglect of negative feedback, whereas values near 0 indicate equal learning from negative and positive feedback.

#### Initial bias.

Although we initialize the Q-values to a fixed value for all stimuli and actions, participants may come in with a preference for pressing action 1 with their index finger, for example (or other biases). To account for subjective biases in action choices and allow for better estimation of other parameters, we use the first choice made by a participant for each stimulus as a potential marker of this bias, and introduce an initial bias parameter *init* that boosts the initial value of this first, prior to the first learning update. Specifically, the initial bias update is Q_0_(s,a_first_(s)) = 1/n_A_+*init\**(1-1/n_A_).

#### Additional mechanisms.

We test two additional mechanisms to attempt to provide better fit within the “RL only” family of models. The first mechanism assumes that learning rates may be different for each set size. The second mechanism assumes that two sets of RL values are learned in parallel and independently, with independent learning rates: one process controls learning phase policy; the other controls testing phase policy only. The best-fitting model in the “RL only” family of models includes all mechanisms described above, for a total of 9 parameters (softmax *β*, 4 learning rates [*α*_learn_(3), *α*_learn_(6), *α*_test_(3), *α*_test_(6)], decay, *pers*, undirected noise *ε*, initial bias). This best model (***RLs***), in addition to the simplest RL model (for baseline), are simulated in Fig. 3 for predictions. The RL model predicts no effect of phase or set size on performance (top row of Fig. 3: the two model learning curves are overlapping); the RLs model’s predictions are dependent on the values of the 4 learning rate parameters, but the model is flexible enough to capture opposite effects of set size in learning and test phases (second row of Fig. 3).

**Figure 3:**
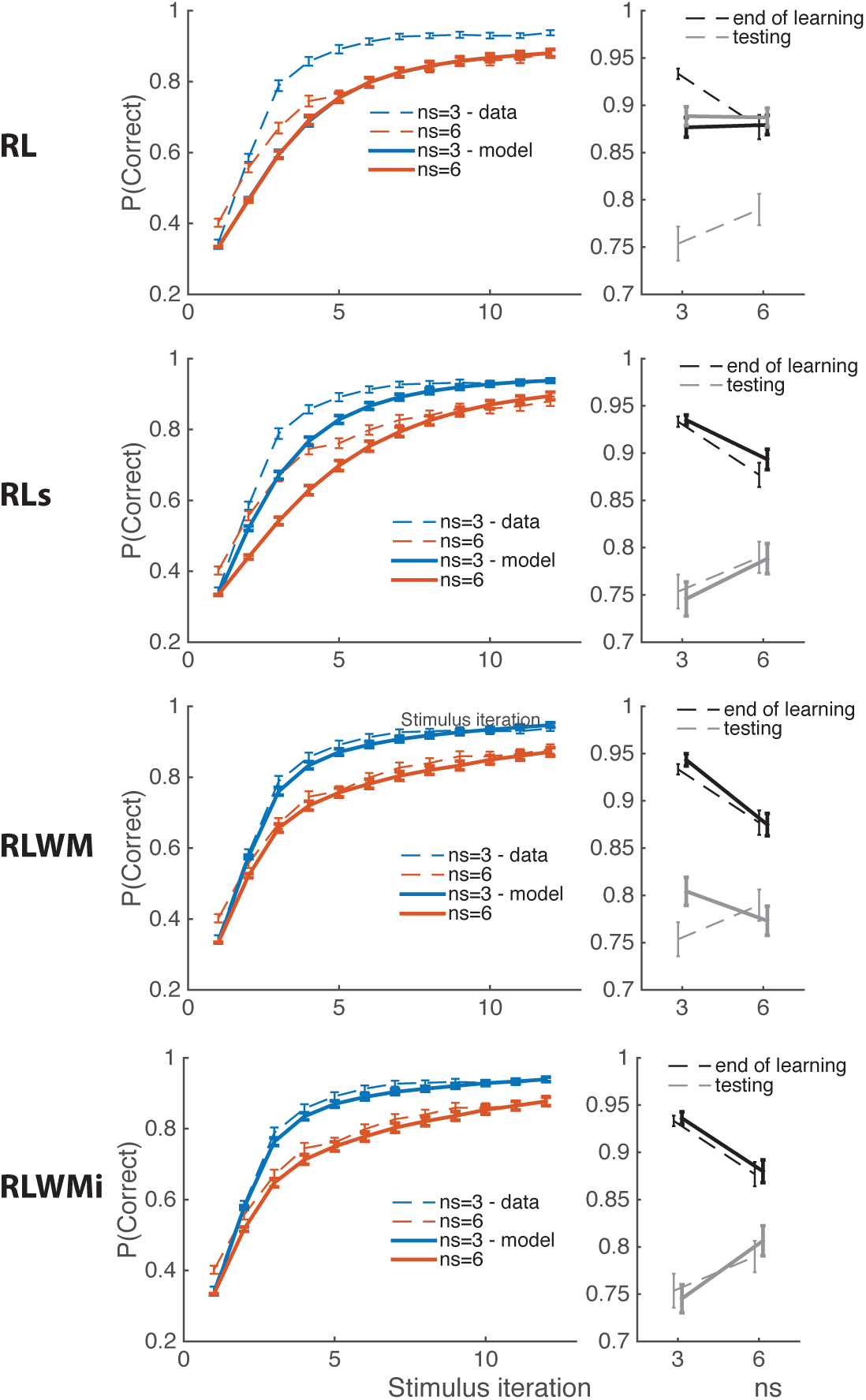
Model Predictions. Model simulations with fit parameters. **Left)** Proportion of correct trials as a function of stimulus iteration number, for set sizes *ns = 3/6*. **Right)** Proportion of correct trials at asymptotic learning performance (last 5 iterations of a stimulus – black line) and during test phase – grey line). Simulations were run with best fit parameters, 100 times per subject. Models are RL (classic two-parameter RL), RLs (best model in the RL family, including two independent Q-tables for learning and testing, and 2 learning rates per set size per learning process), RLWM (with independent working memory and reinforcement learning modules), and RWMi (with interacting modules). Dotted lines show participants’ behavior; full lines model simulations. Note that the simulated learning curves for the RL model overlap, showing no effect of set-size.

### RL+WM independent models

Models in the RLWM family include separate RL and WM modules, and assume that learning phase choice is a mixture of the RL and WM policy, while testing phase choice follows RL policy.

The RL module of the RLWM models is a classic RL model as described above, characterized by parameters α and β, with the additional mechanisms of initial bias and perseveration.

The WM module stores information about weights between stimuli and actions, W(s,a), which are initialized similarly to RL Q-values. To model fast storage of information, we assume perfect retention of the previous trial’s information, such that W_t+1_(s_t_,a_t_) = r_t_. To model delay-sensitive aspects of working memory (where active maintenance is increasingly likely to fail with intervening time and other stimuli), we assume that WM weights decay at each trial according to W_t+1_**←** W_t_ + ϕ (W_0_−W_t_). The WM policy uses the W(s,a) weights in a softmax with added undirected noise, using the same noise parameters as the RL module.

Contrary to previous published versions of this model (Collins et al., 2014; Collins, Ciullo, et al., 2017; Collins & Frank, 2012), we cannot estimate WM capacity directly in this protocol, because we only sample set sizes 3 and 6. Thus, we model the limitations of WM involvement in choice with a fixed set-size-dependent mixture parameter *η*_ns_ for the overall choice policy:

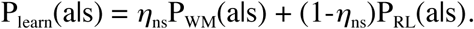

For the testing phase, we assume that P_test_(a|s) = P_RL_(a|s).

The best model in this family thus includes 8 parameters (softmax *β*, RL learning rate *α*, WM decay *ϕ, pers*, undirected noise *ε*, initial bias, two mixture parameters *η*_3_ and *η*_6_). This model predicts identical effects of set size in the learning and testing phases, with worse performance in lower set sizes (Fig. 3, third row).

### RL+WM interacting models

This family of models is identical to the previous one, with the exception that the WM module influences the RL computations. The RL module update is still assumed to follow *Q_t+1_(s,a)* = *Q_t_(s,a) + α x δ_t_*,. However, WM contributes cooperatively to the computation of the reward prediction error *δ_t_* by contributing to the expectation in proportion to WM’s involvement in choice:

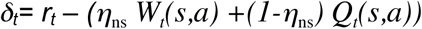

The best-fitting model in this family, *RLWMi*, has the same parameters as the no-interaction RL+WM models. This model predicts opposite effects of set size in the learning and test phases, with worse performance in lower set sizes (Fig. 3, third row). We tested a variant (*RLWMii*) that assumes independent *η*_ns_ parameters for policy mixture and interaction mechanisms; it provided a significantly worse fit (Fig. 4). We also explored a competitive interaction model, whereby WM inhibited RL computations by decreasing learning rate. This model fit equally well as RLWMi, but cannot account for previous EEG findings (Collins & Frank, submitted; see discussion), thus we only report here the cooperative interaction.

**Figure 4:**
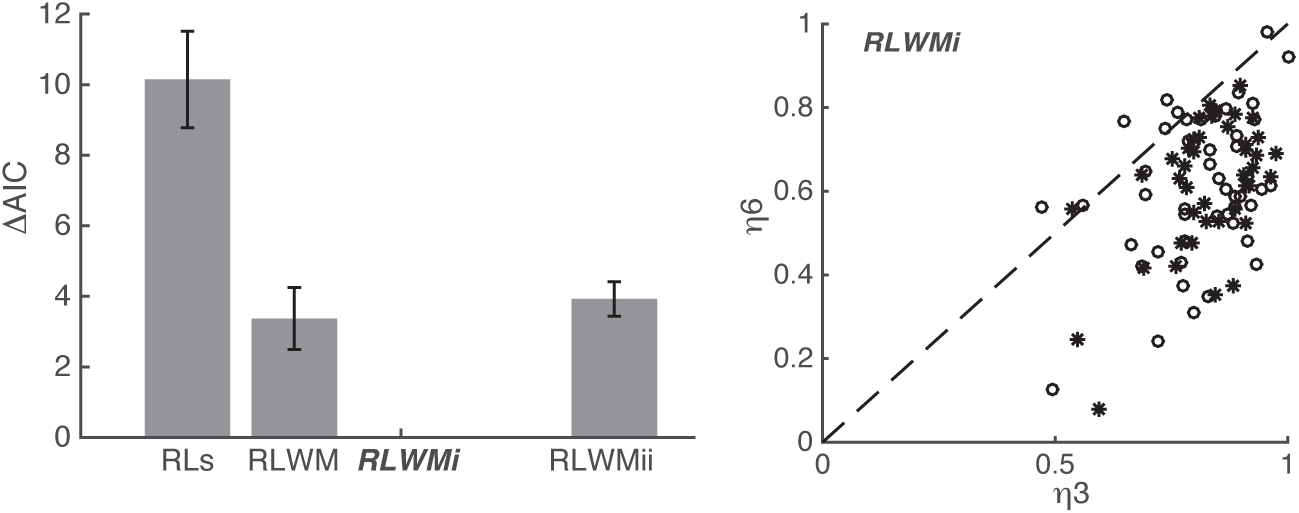
Model fitting. **Left:** Difference in AIC between different models and best model RLWMi. The RLs model is a flexible model including only RL mechanisms; the RLWM model includes an RL module and a working memory module that are independent, but compete for choice. RLWMi additionally includes an interaction, whereby WM contributes to RL computations. RLWMii allows separate weighting for WM’s contribution to choice and to RL computations. Error bars indicate standard error of the mean. **Right:** Mixture weights η indicate contribution of WM to choice and RL computations in both set sizes for all subjects (initial experiment: open circles; replication: stars). As expected under the hypothesis that the WM module represents working memory, we observe consistently lower contribution of WM in high than in low set sizes (*η*_6_ < *η*_3_).

### Model fitting and validation

We used the Matlab constrained optimization function fmincon to fit parameters (the Mathworks Inc., Natick, Massachusetts, USA). This was iterated with 20 randomly chosen starting points, to increase the likelihood of finding a global rather than local optimum. All parameters were fit with constraints [0 1], except the softmax parameter β, which was constrained to [0 100].

We used the Akaike Information Criterion to penalize model complexity (AIC; Akaike, 1974). Results were identical when using Bayesian Information Criterion (BIC; Schwarz 1978), which was used to compute exceedance probability. Comparing the RL-only models, independent RLWM, interacting RLWMi, and double interacting RLWMii, the simple interacting RLWMi model was strongly favored with exceedance probability of *1-5e^−6^* over the whole group (Stephan, Penny, Daunizeau, Moran, & Friston, 2009).

Model selection alone is insufficient to assess whether the best fitting model sufficiently captures the data. To test whether models capture the key features of the behavior (e.g., learning curves), we simulated each model with fit parameters for each subject, with 100 repetitions per subject then averaged to represent this subject’s contribution (Fig. 3).

## Results

Results from the learning phase replicated previous experiments (Collins & Frank, 2012)(Fig. 2A, 2D top). Specifically, we found that participants learned to select correct actions in both set sizes (Fig. 2A top), but were slower to learn in blocks with a set size of *ns* = *6*. This was characterized in a logistic regression analysis which identified two main contributions to learning. First, reinforcement learning was characterized by a positive sensitivity to reward history (number of previous correct trials; sign test p < 10e-4): the more previous correct choices they had, the more likely participants were to pick the correct choice on the current trial. Second, working memory was characterized by a negative effect of set size and delay on performance since the last correct iteration of the current stimulus (both p’s < 10e-4): participants were less likely to pick the correct actions under higher load, or if there had been more intervening trials since they last chose correctly. Furthermore, we identified a practice effect, such that performance was higher in later blocks (sign test, p = 0.01). These results were also paralleled in reaction times (Fig. 2A bottom), such that factors facilitating correct choice (reward history and block) also lead to faster choices (p < 10e-4 and p = 0.002, respectively), and factors making choice harder lead to slower choices (set size and delay; both p’s < 10e-4). At its asymptote, performance was lower in high than low set sizes (t(84) = 6.8, p < 0.0001; 66/85 participants: p < 0.0001). This effect was coupled with higher reaction times (t(84) = 26.9, p <0.0001; 85/85 participants: p < 0.0001).

Model simulations showed that a classic reinforcement learning model could not account for the learning phase effects (Fig. 3 RL), and that while more complex single process reinforcement learning models can capture a qualitative effect of set size with set size dependent learning rates, they do not quantitatively capture the learning phase dynamics (Fig. 3 RLs). In contrast, we replicate our previous finding that RLWM models can capture learning phase dynamics well (Fig. 3 RLWM, RLWMi).

We next analyzed testing phase performance as a function of the set size in which an item had been learned. As expected from our hypothesis that learning relies on RL and WM, but testing on RL only, we found overall worse performance (t(84)=10.5, p<10e-4) in the testing phase compared to asymptotic performance in the learning phase, as well as faster reaction times (t(84)=21.1, p<10e-4). Furthermore, our theory assumes that learning in high set size blocks relies more on RL than in low set size blocks, and that testing phase relies only on RL. Thus, we predicted that individual differences in testing phase behavior should be better predicted by individual differences in asymptotic learning phase performance in high than low set size blocks, and that this should be true for stimuli learned in either set size. Unsurprisingly, a multiple regression analysis with both asymptotic set size 3 and 6 learning phase performance showed that set size 6, but not set size 3, was significantly predictive of set size 6 test performance (t(84) = 5.06, p < 10e-4 for ns = 6; t(84) = 1.8, p > .05 for ns = 3). Surprisingly, but confirming our prediction, we also found that set size 3 testing phase performance was predicted by set size 6, but not set size 3 asymptotic learning phase performance (t(84) = 4.5, p = 0.0000 for ns = 6; t(84) = −0.02; p = 0.98 for ns = 3). These results support our hypothesis that testing phase performance relies on RL, which is more expressed during learning of set size 6 than 3.

Next, we investigated testing phase performance as a function of set size of the block during which the stimulus was learned. It is important to remember that if testing phase performance reflected the history of choices and rewards during learning, participants should be better able to select correct actions for stimuli in low set sizes rather than in high set sizes (following their asymptotic learning performance). Instead, we found that participants were significantly better at selecting the appropriate action for items they learned in high set sizes (Fig. 2B t(84) = 3.8, p = 0.0003; 58/85 participants: sign test p = 0.001), indicating a counterintuitive robustness of learning under high load. This was also visible in reaction times, which were faster for items learned under high load (Fig. 2B t(84) = 3.5, p = 0.0008; 57/85 participants: p = 0.002).

We confirmed these results with multiple regression analyses, using learning block, learning set size, and asymptotic average rewards as predictors. Results showed that learning block and reward history accounted for significant variance in testing phase performance and RTs (Fig. 2D; sign tests p’s < 10e-4), with effects in the same direction as learning phase performance and RTs, reflecting variance in the learning process. However, set size predicted variance in the testing phase in the opposite direction from in the learning phase (Fig. 2D; better choices, p < 10e-4, faster RTs, p = 2.10e-4). Together, these results confirm our prediction that despite apparent successful learning in low set sizes, stimulus-action associations were formed in a way that was stronger in high set sizes, as observed in testing phase performance.

Next, we sought to confirm that these effects were not due to differences in reward history. First, we limited testing phase analysis to items where the participant chose correctly for all last 6 iterations (thus reaching 100% asymptotic performance in both set sizes). Within this subset, we again observed better test performance in high set size stimuli, with a stronger effect (p < 10-4, t(84) = 7.6). Second, we asked whether higher performance in set size 6 might be accounted for by participants better avoiding choices that were unrewarded during learning, which would have been experienced more in set size 6 than set size 3. However, we found that the types of errors made by participants in both set sizes were not different (t(84) = 0.01,p = 0.99), and significantly suboptimal (one-sample t-test vs. chance both t’s > 12, proportion of optimal error trials 27%; see methods). Thus superior performance in high set sizes was not due to a better avoidance of incorrect actions due to higher sampling of errors during learning. Indeed, RLWM models with no interaction between RL and WM predicted the *opposite* effect of set size in the testing phase (Fig. 3 RLWM).

Instead, we hypothesized that the reversal in the effect of set size on performance between learning and testing was indicative of WM interfering with RL computations during learning. Specifically, we hypothesized that in low set size blocks, which were putatively within working memory capacity, the ability to solve the learning problem with WM would lead to interference with RL computations. We propose a mechanism whereby WM is able not only to store stimulus action associations, but also to predict a correct outcome when this association is used. This prediction is used cooperatively in a mixture with the RL expectation to compute reward prediction error (Fig.1C); and thus when WM is ahead of RL, it decreases positive reward prediction errors, and effectively impedes learning in the RL process (Collins & Frank, submitted). This would then lead to less well-learned associations in low compared to high set sizes. We characterized the contributions of WM and RL to choice and to the reward prediction error with a single mixture parameter for each set size, η_ns_ (see methods for full model description). If this mixture parameter represents contributions of capacity-limited WM, we should observe η_6_<η_3_. This is indeed what we observed with parameters fit on individual subjects’ choices (Fig. 4 right).

We tested this model (RLWMi) against three other main models (as described in the methods). The first alternative (RLs) assumed only RL processes, and was made flexible enough to be able to reproduce all qualitative effects observed empirically. This required assuming that two independent sets of association weights were learned in parallel, one used during learning and the other during testing; and that learning rates were different for both, as well as for different set sizes. The second alternative model (RLWM) was similar to the main model (RLWMi), but without the interaction for learning. The last alternative model (RLWMii), was identical to RLWMi, but offered flexibility in separately parameterizing the contribution of WM to choice and to learning. We fitted participants’ behavior with the three models, and found that RLWMi fit significantly better than the other three models (exceedance probability = 1). Model simulations (Fig. 3) confirmed that RLWMi provided the best qualitative and quantitative account of the data. These results confirm that working memory is needed to account for learning behavior, and that an interference with RL is the best way to account for counterintuitive testing phase effects. It is important to note that letting the contributions of WM to choice and to RL be independent does not capture additional variance in behavior, hinting that these two functions might share a single, coupled mechanism.

To validate the RLWMi model, we simulated it with individual participants’ fit parameters (100 times per participant). Simulations provided a good qualitative and quantitative fit to the choice results in both learning and testing phase. Specifically, the simulations produced learning curves that were very close to participants’ (Fig. 5A), and reproduced the opposite set size effects in asymptotic learning phase and testing phase (Fig. 5B, D, E). We did not capture the practice effect (block; Fig. 5D, E). We also simulated reaction times by assuming a negative linear dependence on the model-determined probability of the choice, such that faster reaction times occurred for choices with higher confidence. Using these simulated reaction times, we reproduced the observed pattern whereby reaction times are faster for low set sizes than high set sizes during learning, but the opposite is true for the testing phase (Fig. 5C). This is because during learning, the WM contribution to choice leads to more confidence in the choices and faster RTs; but the higher WM interference during learning leads to less well-learned Q-values for low set sizes, which translates to slower RTs during testing. It is important to note that the model was fit only to the choices, not RTs, so this is an independent test of the model’s ability to capture empirical data.

## Discussion

Our new experimental protocol allows us to demonstrate a surprising finding: associations learned in a more complex problem end up being learned more robustly, and are better retained in the long-term, than easy to learn associations. This finding provides behavioral evidence for an interaction between working memory and reinforcement learning. Specifically, we showed previously that RL and WM compete for choice during learning (Collins et al., 2014; Collins & Frank, 2012). Here, we show an additional interaction between them whereby working memory impedes RL computations. These new results behaviorally and computationally support our previous observations that WM and RL are not independent modules, but that RL neural signals are weakened by WM use (Collins, Albrecht, et al., 2017; Collins, Ciullo, et al., 2017)(Collins & Frank, submitted). It also strengthens our previous result showing that stimulus value was better learned in high set sizes (Collins, Albrecht, et al., 2017), but offers a more robust computational account for this finding.

Working memory and reinforcement learning implement different trade-offs for learning: WM allows for very fast learning of information that is not durably retained, while RL allows for slow, integrative learning of associations that are robustly stored. Previous models assumed that humans can benefit from the best of both worlds by shifting the weight given to each system as a function of their reliability (Collins et al., 2014; Collins, Ciullo, et al., 2017; Collins & Frank, 2012). Specifically, assuming that the effortless RL process simply occurs independently in the background, we can use WM to the maximum of its reliability for learning, and use RL first as a back-up, but as it becomes more reliable than WM, shift to “automatized” RL behavior only. The results presented here show instead that the trade-off cannot be completely eliminated by carefully arbitrating between RL and WM during learning: the use of WM in easy problems weakens RL learning, and thus leads to faster learning at the cost of long-term retention.

We proposed a computational mechanism by which working memory may impede RL learning. Despite its negative effect on long-term learning, this mechanism can be thought of as cooperative. Indeed, we suggest that when an association is held in working memory, working memory can also contribute to corresponding reward expectations. Our computational mechanism assumes that WM’s expectation is weighted with RL’s expected value, and that this mixture expectation is then used to compute RL’s reward prediction error. Thus, when WM is reliable and learns faster than RL, this mechanism generates lower reward prediction errors in correct trials than an independent RL mechanism would, which in turn leads to a weaker update of associations in the RL system. This mechanism is compatible with our observations in an EEG experiment, where trial-by-trial neural markers of working memory use predicted lower markers of reward prediction errors (Collins & Frank, under review).

We showed that this cooperative interaction accounted for trial-by-trial choices during learning and testing better than models assuming no interactions. However, other interaction mechanisms could also be considered - in particular, a competitive mechanism provides a similar fit to participants’ choices (Collins & Frank, submitted). This mechanism simply assumes that the RL learning rate is decreased, in proportion to WM contributions to choice. This leads to slower RL learning, not due to weakened reward prediction errors as in the cooperative mechanism, but due to a weaker effect of reward prediction learning on updates. Competition and cooperation mechanisms make separable predictions for neural signals: the competition mechanism predicts that RPEs will decrease slower in low set sizes due to the slow learning rate, while the cooperation mechanism predicts that they will decrease faster in low set sizes, due to accurate WM predictions. EEG results confirmed the latter prediction (Collins & Frank, submitted), thus supporting the cooperation mechanism prediction. However, it will be important in future work to show behavioral evidence disambiguating the two mechanisms.

This leaves opens an important question – are there situations in which a cooperative mechanism might be beneficial? In our experimental protocol, it seems suboptimal, as it leads to weakened learning in the long-term. RL algorithms are only guaranteed to converge to true estimates of expected value if the reward prediction error is not tampered with; interference may bias, or slow computations, as is seen here. It is thus puzzling that we might have evolved an interaction mechanism that actively weakens one of our most robust learning systems. Future research will need to determine whether this bias might be normative in more natural environments – for example, it is possible that situations in which learning is required to be very fast are also volatile environments, where we might need to change our behavior quickly. In that case, not having built as strong an association might allow for more flexible behavior. Another hypothesis is that this interaction reflects constraints of neural network implementations, and is thus a side-effect of another normative mechanism: specifically, that the contribution of working memory to choice, which helps learn faster in low set sizes, is not separable from its contributions to the reward expectation (Fig. 1C). While we do not have direct evidence for this hypothesis, our model fitting results provide a clue in its favor: a more flexible model that allowed contributions of WM to choice and RL to be uncoupled did not provide a better fit, and the additional degrees of freedom led to overfitting. Thus, no additional variance was captured by letting choice and learning interactions be separable, pointing to the possibility that they are indeed coupled mechanisms. Future research will need to clarify this.

We focused on how two neuro-cognitive systems, working memory and reinforcement learning, worked together for learning, highlighting the effects of their interactions in a testing phase where only reinforcement learning was used. However, it is likely that one other system – long term memory – contributes to testing performance, and potentially also to learning (Bornstein, Khaw, Shohamy, & Daw, 2017; Bornstein & Norman, 2017). Indeed, others have shown that long-term memory encoding may compete for resources with reinforcement learning processes (R. A. Poldrack & Packard, 2003; Wimmer, Braun, Daw, & Shohamy, 2014), and other interactions may be possible (for a review, see (Gershman & Daw, 2017)). Here, we observed in the testing phase that participants performed better for associations learned closer to the testing phase, an effect reminiscent of the well-documented recency effect in long term memory recall (Sederberg, Howard, & Kahana, 2008). Our model currently does not capture this effect, or other potential contributions of long term memory. While including it is beyond the scope of this work, it will be important in future research to investigate how this third mechanism interacts with RL and WM for learning.

In summary, our results show a counter-intuitive but robust (and replicable) finding – that while learning under high load is slower and more effortful, it actually allows for better long-term learning and retention. This appears to be due to the fact that faster learning under low load cuts the corner with working memory, and by doing so undermines more robust encoding of associations via reinforcement learning. Our findings highlight complex interactions between multiple learning systems not only at the level of decisions, but also at the level of learning computations. When learners arbitrate in favor of fast and efficient working memory use over RL in simple situations, they simultaneously undermine the computations from this slower but more robust system, leading to worse long-term performance. This result may have important implication in numerous domains, as learning is an important part of our daily lives – for example in educational settings. Understanding how multiple systems work together for learning is also a crucial step to identifying the causes of learning dysfunction in many clinical populations, and thus better targeting treatments.

## Acknowledgements

I thank Nora Harhen and Sarah Master for data collection.

